# iPS-CNM: an iPSC collection generated by base editing for centronuclear myopathy

**DOI:** 10.64898/2026.09.15.751756

**Authors:** Insa Staecker, Claudia Pommerenke, Amy Ellinghaus, Ida Steinkirchinger, Vivien Hauer, Hannah Kallnischkies, Maren Kaufmann, Lothar Gröbe, Silke Fähnrich, Josephine Haake, Laura Steenpass, Haicui Wang

## Abstract

Centronuclear myopathy (CNM) is a rare form of inherited diseases often caused by single base mutations. Modelling CNM is challenging due to the diversity of CNM mutations and genetic background of individual patient. To address this, we used base editing to introduce CNM mutations into an induced pluripotent stem cell (iPSC) line from a healthy donor, generating a collection of iPSC lines (iPS-CNM) carrying distinct CNM mutations. We found that the efficiency of base editing depended critically on selecting base editor (BE) variants and the target sequences. Optimization using different BE variant and sgRNA pair was required for each target site. Moreover, whole genome sequencing (WGS) was performed to confirm on-targets and detect off-targets in order to select iPSC clones for the collection. Our findings highlight the feasibility of base editing for generating an iPSC collection from one parental iPSC line combined with thorough evaluation of rare off-targets using WGS.

## Introduction

Centronuclear myopathy (CNM) is a rare form of inherited diseases. So far, mutations in the following genes have been reported to cause CNM: recessive X–linked mutations in *MTM1*^1^; autosomal dominant (AD) mutations in *DNM2*^2^ or *BIN1*^3^; autosomal recessive (AR) mutations in *BIN1*^4^, *RYR1*^5^*, TTN*^6^ or *SPEG*^7, 8^. The diversity of mutations carried by individual patients is reflected by the broad range of clinical manifestations, which brings challenges in making a genotype–phenotype correlation and developing personalized treatment.

Base editors (BEs) which conduct direct conversion of single nucleotide without making double–strand DNA breaks (DSBs) hold high potential in disease modeling or treatment of genetic diseases.^9, 10^ Among CNM mutations, the majority are single nucleotide variants (SNVs) such as missense, nonsense or splice site mutations. Taking the severe X-linked CNM as an example, approximately half of the *MTM1* mutations identified to date (supplementary Fig. S1) can be either corrected by BEs in patient genomes to explore potential therapeutic strategies, or introduced by BEs to healthy genetic backgrounds for disease modeling.

BEs are composed of a Cas9 coupled with a deaminase enzyme. The classical cytosine base editor (CBE) uses a cytidine deaminase to convert C-to-T (G-to-A)^11^ and adenine base editor (ABE) uses an adenosine deaminase to convert A-to-G (T-to-C)^12^. Further engineered BEs allow precise base editing across all four nucleotides, substantially expanding the editing range of BEs.^13, 14, 15^ Like other CRISPR tools, BEs require a single guide RNA (sgRNA) to bind to the target sequence and a Protospacer Adjacent Motif (PAM) following the target sequence to mark proper target sites. Their maximum editing efficiency can be achieved by positioning the targeted nucleotide within the editing window. Earlier versions of BEs using NGG PAM limit target site recognition due to sequence constraint. Further engineered Cas9 variants with relaxed PAM such as SpG (NGN PAM) or SpRY (NRN/NYN PAM: R=A/G, Y=T/C) removed such constraint, yet they also resulted in relatively lower editing efficiency and a higher risk of off-target effects.^16^

Generation of an isogenic, induced pluripotent stem cell (iPSC) collection using gene editing is increasingly utilized to establish disease models, as iPSC can turn into almost any cell type in the human body^17, 18^. Although BEs are designed to avoid DSBs, recent research shows that they can still induce unintended genomic alterations, including structural variations (SVs) such as large deletions, truncations, and aneuploidy, small Indels (insertions and deletions) and SNVs.^19,20^ Theoretically BEs can create a collection of isogenic iPSC lines by introducing SNVs of one disease type to a single parental iPSC line, to investigate the mutation effects within a presumably identical genetic background. However, any unintended genomic alterations introduced to iPSC lines during base editing might question the observed single mutation effect for monogenic diseases such as CNM.

In this work, we explore the editing efficiency and safety using BEs to generate an iPSC collection consisting of cell lines carrying individual CNM mutation. We attempted to introduce 10 individual mutations of 4 CNM genes to one iPSC line from a healthy donor through variable combination of BE variants and sgRNA sequences. For generated iPSC CNM lines, we detected the on-targets and off-targets using whole genome sequencing (WGS) and the single-donor background facilitates the extraction of this information through direct comparison of the edited lines with the parental line. We demonstrated that BEs enabled the generation of iPSC lines carrying desired CNM mutations from one donor background at appreciable efficiency and safety. Here, we present a summary of our experience on how to achieve on-target editing by adjusting the editing strategy and how to identify high risk off-targets using WGS for iPSC lines.

## Methods

### Base editing experimental design

CNM mutations were selected to be introduced by BEs to one iPSC line (supplementary Table S1), covering different inheritance types and disease severity from 4 CNM genes. The targeted nucleotide was placed within the editing window of BE. Most selected conversions require CBE4max and its variants, which have an efficient deamination window typically from positions 4 to 8 within the protospacer, counting the end distal to the PAM as position 1 in the sgRNA sequence^16^, except one requires an ABE with editing window also from position 4 to 8^21^. In addition, any potential bystander editing targets were excluded from the editing window if possible. And NGG or NGN PAM has higher preference than NRN or NYN PAM for BE selection. The sgRNAs with predicted high specificity and less off-targets were selected using the web tool CRISPOR.^22^

### Cell culture, cell thawing and cryopreservation

The iPSC line RBi001-A purchased from EBiSC - European Bank for induced pluripotent Stem Cells was selected as it is from a male healthy donor which is suitable to model X-linked mutations as well as autosomal mutations. Cells were cultivated at 37°C and 5% CO^2^ in 6-, 12-, 24- or 96-well cell culture Nunclon Delta plate (Thermo Fisher Scientific), coated with hESC-grade Matrigel (Corning) and maintained in iPS-Brew medium (Miltenyi Biotec). Cells were passaged using 0.5 mM EDTA (Invitrogen) when cultures reached 80% confluency. Cells were thawed in iPS-Brew supplemented with 10 µM Y-27632 (Stem Cell Technologies), followed by medium switch to iPS-Brew medium after 24 hours. Single iPSCs were achieved through dissociation with Accutase (Gibco) and cryopreserved with serum-free cryopreservation medium Bambanker (Nippon Genetics) either for short time at -80°C or long time in liquid nitrogen tank.

### Vector construction

The sgRNA plasmids were generated by replacing the existing sgRNA sequence with designed CNM sgRNA sequences in pAT15415-BEAR-GFP-target-mCherry plasmid (Addgene #162995, from Ervin Welker lab^23^), using Q5 Site-Directed Mutagenesis Kit (NEB). See Supplementary Table S2 for the full list of oligonucleotides used to generate sgRNA constructs.

The BE plasmids were purchased from Addgene: pCMV_BE4max_P2A_GFP, Addgene # 112099 from David Liu lab^24^; pCAG-CBE4max-SpG-P2A-EGFP, Addgene #139998 and pCAG-CBE4max-SpRY-P2A-EGFP, Addgene #139999, both are from Benjamin Kleinstiver lab^16^; pCMV-T7-ABE8e-SpRY-HF1-P2A-EGFP, Addgene #197507 from Benjamin Kleinstiver lab^9^. The transfection grade plasmid DNA was prepared using the QIAGEN Plasmid Midi or Maxi Kit.

The schemes of representative BEs and sgRNA plasmid maps can be seen in supplementary Fig. S2.

### iPSC transfection with BE and sgRNA vectors

The transfection was performed as previously described.^25^ Briefly, the iPSCs were seeded on 6-well plate coated with Matrigel at a density of 3.1×10^4^ cells/cm^2^ in iPS-Brew medium with 10 µM Y-27632. After 24 hours, cells were switched to fresh iPS-Brew medium and transfected with 1 µg BE plasmid and 1 µg sgRNA plasmid using Lipofectamine 3000 Reagent (Thermo Fisher Scientific) following manufacturer’s instructions. The medium was replaced approximately 24 hours after transfection. Cells were sorted 48 hours after transfection using a FACSAria II or FACSymphony S6 cell sorter (BD Biosciences) for EGFP and mCherry double-positive cells. Sorted iPSCs were maintained in iPS-Brew medium with 10 µM Y-27632, 1X CloneR™2 (StemCell Technologies), 100 μg/mL Primocin (InvivoGen) for 1-3 days until small iPSC colony formed, followed by expansion in iPS-Brew medium with 100 μg/mL Primocin for another 3-5 days.

### Single clone isolation of iPSCs for expansion

The iPSCs were passaged using Accutase and seeded at low densities (around 50 and 100 cells/cm^2^) for single colony formation in 6-well Matrigel-coated plates and maintained in iPS-Brew with 10 µM Y-27632 until small colonies formed. The iPSC colonies were further expanded for 5-7 days until middle size (100-200 µm), single iPSC colonies were picked (>30 for each editing) into Matrigel-coated 96-well plate in iPS-Brew medium for expansion and subsequent genotyping. The rest cells were pelleted as pooled cells for Sanger sequencing to estimate the overall editing efficiency for some cases.

Single edited clones verified after genotyping were propagated to passage at least 3-5 times. The authenticated and mycoplasma-free clones further went for banking, characterization regarding iPSC morphology, iPSC pluripotency as previously described^26^, and WGS.

### Sanger sequencing to screen for edited clones

Single clones were expanded from 96-well plate into 2 wells in 24-well plate, with one well for genotyping and 1 well for further expansion as clone preservation.

Genotyping was carried out as described previously.^25^ Briefly, genomic DNA (gDNA) was isolated with Agencourt AMPure XP beads (Beckman Coulter) according to manufacturer’s instructions. PCR fragment longer than 200 bp containing the target sequence was amplified using OneTaq DNA Polymerase (NEB) and purified by QIAquick PCR Purification Kit (Qiagen), then sent to Sanger sequencing (Microsynth). Sequence chromatograms were analyzed with EditR^27^ (version 1.0.10). Primers used for sequencing were listed in Supplementary Table S3.

### Whole genome sequencing (WGS) and data analysis

The gDNA of selected iPSC CNM lines (Supplementary Table S4) and unedited parental control line was isolated with QIAamp DNA Blood Mini Kit (Qiagen). Library preparation was performed by Novogene using human whole genome library. The library was sequenced on Illumina NovaSeq X Plus instrument with a read length of 2 x 150bp and 30 x coverage.

After preprocessing via fastp (0.23.4)^28^, trimmed reads were aligned to the human reference GENCODE (hg38, v42)^29^ via bwa (bwa-0.7.17-r1188)^30^. Mapped reads were sorted, whereas discordant and split reads were separated by samtools (samtools 1.18, htslib 1.17)^31^ in order to prepare mapped reads for structural variant (SV) detection. Different SV caller Control-FREEC (v11.6)^32^, DELLY (v1.3.3)^33^, LUMPY (v0.2.13)^34^, and Manta (1.6.0)^35^ were applied to search for genomic rearrangements.

Sorted mapped reads served also for SNV and small Indel detection using Mutect2^36^ following the pipeline described elsewhere^37^ except for starting with bwa instead of star aligned reads and the last filtering steps protein coding and frequency. The possible damaging SNV/Indels including on-targets were filtered from SNV/Indels of coding exons and splice site/regions, by excluding the ones with SIFT score ≥ 0.1 and PolyPhen benign, but keeping those without SIFT score and PolyPhen prediction for analysis to avoid over filtering. The functional interaction network of perturbed proteins by refined possible damaging SNVs/Indels was generated using STRING^38^. Off-targets analysis were conducted by comparing SNV/Indels (coding exons, 3‘UTR, 5‘UTR, splice sites/regions) from WGS detection with predicted exon off-targets (maximum 4 mismatches) by CRISPOR^22^ and CCTop tool^39^. The shared off-target genes were nominated first following by mapping the predicted off-targets in genomic loci using IGV browser. The positive mapped off-targets were considered clone-specific off-targets.

The WGS data has been deposited at ArrayExpress: E-MTAB-17661.

### iPSC Pluripotency by immunofluorescence staining (IF)

Cells were fixed with 4% paraformaldehyde (PFA) for 15 min at room temperature. Primary antibody was diluted 1:200 in in blocking buffer (5% milk powder, 0.4% Triton-X-100 in PBS) and incubated for 1 hour at room temperature. After a three-times PBS wash the secondary antibodies were applied, diluted 1:1,000 in blocking buffer for 30 min. After PBS wash, the cells were incubated for 20 min with DAPI solution (5 µg/ml in PBS, Sigma Aldrich) and investigated with a Zeiss Axiovert A1 microscope.

Primary antibodies (Cell signaling): anti-Nanog (rabbit polyclonal), anti-Oct-4A (rabbit polyclonal), anti-Sox2 (rabbit polyclonal). Secondary antibodies: anti-Rabbit IgG Alexa Fluor 488 (Cell signaling) or Alexa Fluor® 594 goat anti-rabbit IgG (H + L) (Thermo Fischer Scientific).

### iPSC Pluripotency by flow cytometry

Single cells dissociated with Accutase were incubated in PBS with respective antibodies (dilution 1:25) for 30 min at 4°C and washed three times by PBS. Cells were analyzed in a 96-well plate at a Beckman Coulter CytoflexS flow cytometer. Results were plotted using CytExpert software. Antibodies (Miltenyi Biotec): anti-SSEA-1, PE; anti-SSEA-4, PE; anti-TRA-1-60, PE; anti-TRA-1-81, PE; anti-IgG1, PE.

### Pluripotency analysis: trilineage differentiation

Differentiation was achieved using StemMACS^TM^ Trilineage Differentiation Kit (Miltenyi Biotec) according to manufacturer’s instruction. Single cells dissociated with Accutase were seeded at densities of 4.3×10^4^, 7.1×10^4^ and 5.7×10^4^ cells/cm^2^ for meso-, endo- and ectoderm, respectively.

Staining for trilineage differentiation markers was performed as described by the manufacturer, antibodies were diluted 1:50. A 4% PFA solution was used for fixation and 0.1% Triton-X-100 in PBS for permeabilization. Cells were analyzed in a 96-well plate at a Beckman Coulter CytoflexS flow cytometer. Antibodies (Miltenyi Biotec): anti-CD140b, APC; anti-CD144, FITC; anti-CD184, APC; anti-Sox17, VioB515; anti-PAX-6, APC; anti-Sox2, FITC.

## Results

### Selection of CNM mutations for iPSC line generation

Among mutations of several CNM genes^1–5^, we selected 10 CNM SNVs covering different inheritance patterns including *MTM1* X-linked nonsense and missense mutations, *DNM2* AD missense mutations, *BIN1* AD and AR missense mutations, and *RYR1* AR missense mutations, which also associated with different disease severity (Supplementary Table S1). To assess the single mutation impact in the context of single donor background, we introduced each mutation using BEs to one iPSC line from a male healthy donor which is suitable to model not only X-linked CNM but also other AD and AR CNM. We identified edited clones with heterozygous and homozygous genotypes for each CNM mutation from total analyzed clones (Fig. 1A, Table 1), to assess the editing ratio and editing efficiency from each base editing.

**FIG 1.**
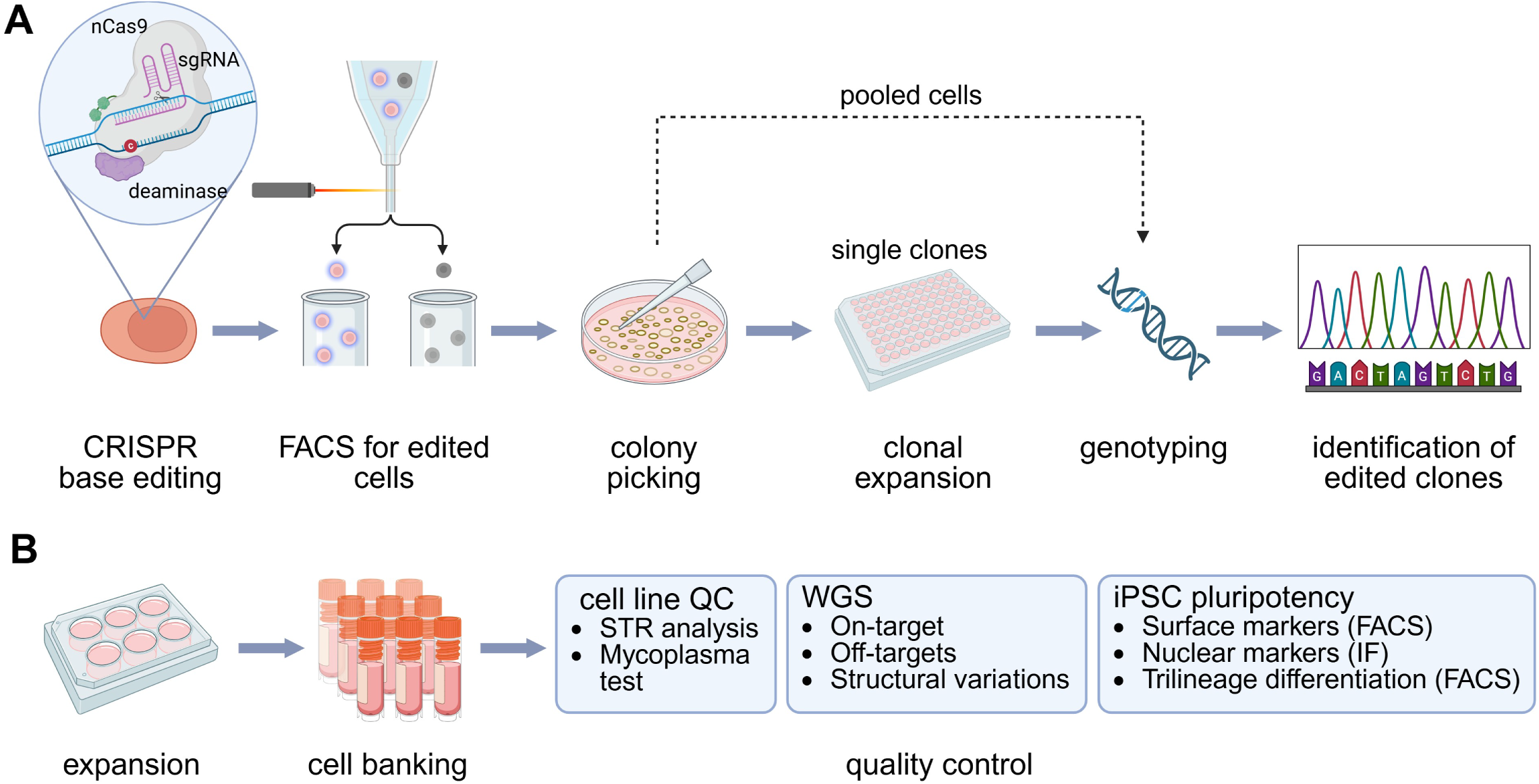
Base editing workflow to generate the iPS-CNM collection. (A) Base editing involved the introduction of CNM mutations into the iPSC line using BEs followed by fluorescent-activated cell sorting (FACS) to isolate double fluorescent positive cells expressing BE and sgRNA. BEs used in this study consist of a Cas9 nickase (nCas9) and a deaminase expressed along with an EGFP reporter from one vector, and the co-delivered sgRNA was expressed along with a mCherry reporter from another vector (supplementary Fig. S2). Single colonies were picked and genotyping was performed to identify edited clones carrying the intended mutation using Sanger sequencing. Genotyping for pooled cells was used to preliminarily assess the editing efficiency. (B) Identified positive clones were further expanded, characterized according to standard cell line QC, iPSC pluripotency QC, and genetic QC using WGS. This figure was created in BioRender (https://BioRender.com/j0pmbo6).

**Table 1.**
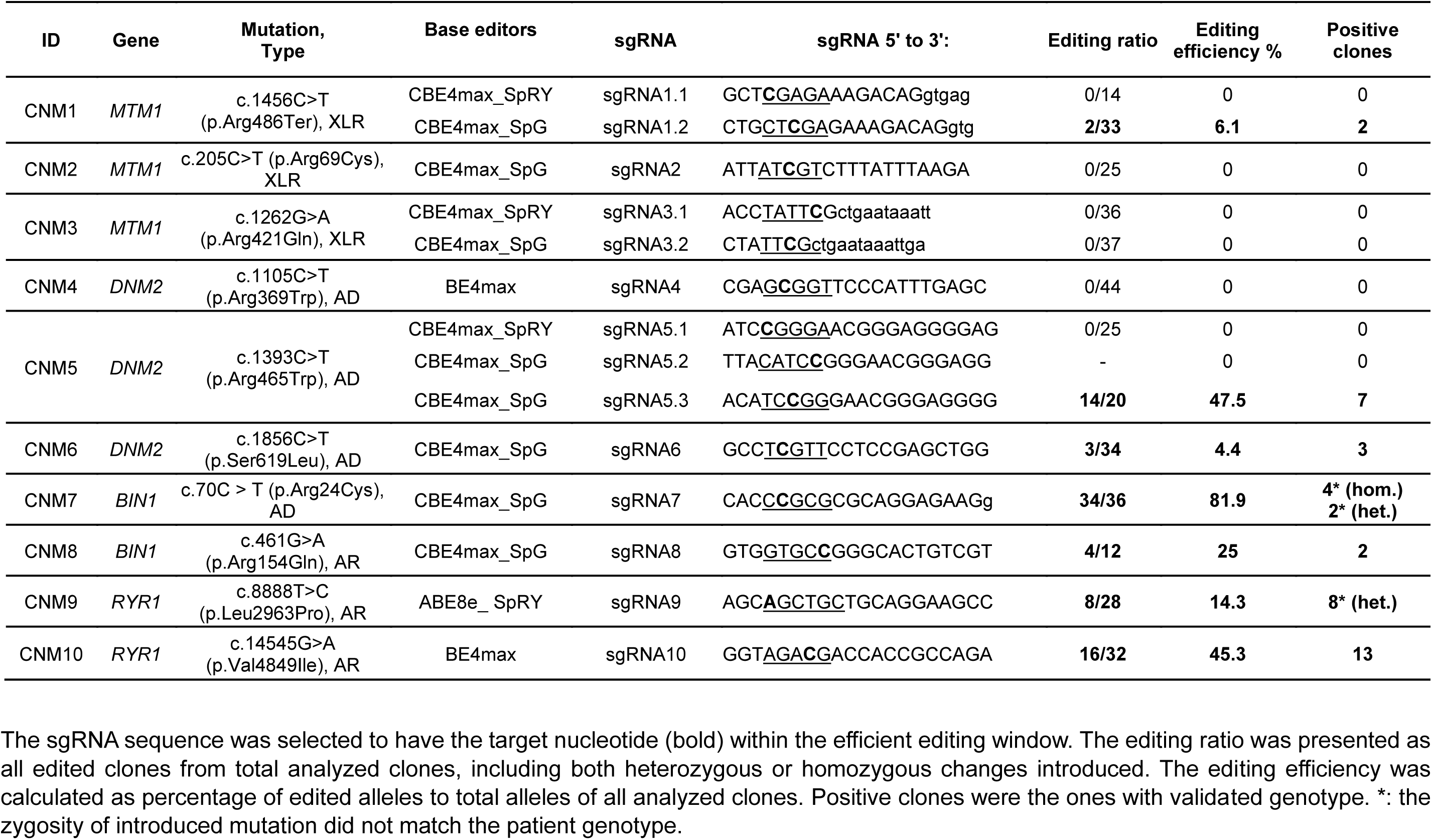
Summary of base editing results.

To generate a collection of iPSC lines for CNM, we selected 1-2 edited clones for each CNM mutation per iPSC line for large banking and iPSC quality control (QC), followed by genetic QC using WGS to validate the on-targets and to identify off-targets or any structural variants (SV) (Fig. 1B). In total 8 iPSC clones from 6 CNM lines were processed in reference to unedited parental cells (Supplementary Table S4) for WGS analysis, aiming to distinguish novel variants introduced by base editing from the pre-existing variants of the parental iPSC line.

### Optimized base editing strategy for high on-target activity

We initially attempted to introduce the desired mutation by placing the targeted nucleotide within the editing window of BE and shifting other nucleotides of the same type outside of the editing window as established previously^25^. Taking the CNM-5 *DNM2* c.1393C>T as example, the first editing was carried out with CBE4max_SpRY and a sgRNA5.1 which has only one C as the target nucleotide within the editing window (Fig. 2A) but predicted to be low specificity and high off-target potential (Supplementary Table S1). This editing strategy resulted in no detectable editing (Fig. 2B).

**FIG 2.**
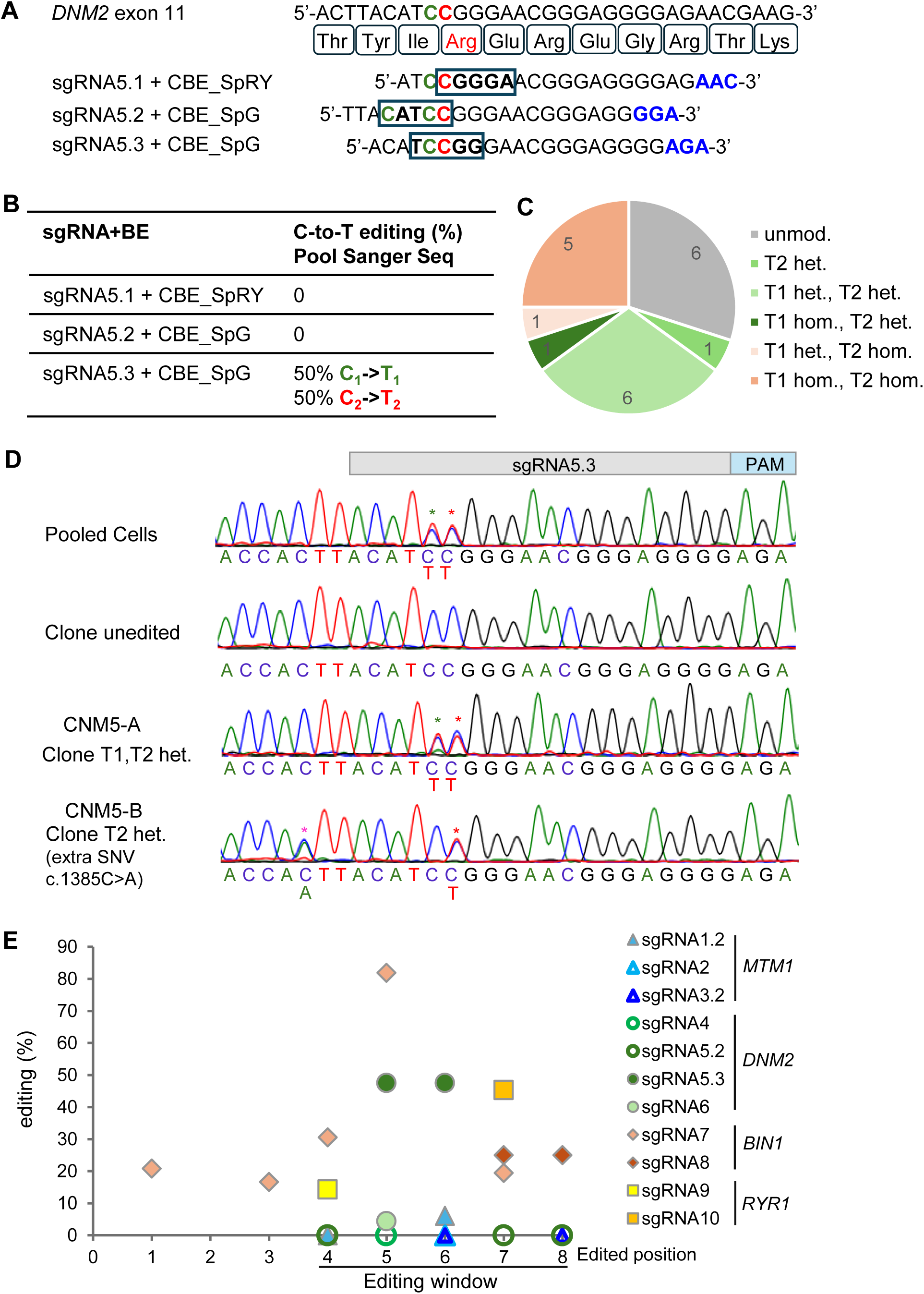
Editing strategy for iPSC CNM5. (A) Introduction of *DNM2* c.1393C>T (p. Arg465Trp) autosomal dominant mutation to iPSCs using different BEs and sgRNAs. The target C was marked in red and the nucleotides of potential bystander editing were marked in green which will not cause protein sequence change in this case. PAM sequence was in blue and editing window was labeled in black frame. (B) Sanger sequencing results of pooled BE and sgRNA expressing cells after sorting. (C) The number of edited clones with different genotypes in total analyzed clones (n=20). Unmod. = unmodified, hom. = homozygous, het. = heterozygous. (D) Sanger sequencing result for pooled cells and representative single clones. Binding site of sgRNA and related PAM are indicated on top. The red * marked the intended changes, and green * marked the synonymous substitution, and the magenta * marked one unintended mutation introduced during base editing or iPSC expansion. (E) Editing efficiency summarized from total analyzed clones. The editing efficiency was calculated as percentage of edited alleles to total alleles of all analyzed clones (Table 1).

Further optimization by adjusting the positions of the target C within the editing window allowed editing with higher specific BE such as CBE4max_SpG^16^ and sgRNAs of predicted higher specificity (sg5.2 and 5.3) despite synonymous bystander editing might occur due to other C in the editing window (Fig. 2A and supplementary Table S1). An improved editing was detected with a roughly 50% c.1393 C-to-T editing and also 50% bystander editing of c.1392 C-to-T from pooled positively transfected iPSCs, using CBE4max_SpG and sgRNA 5.3 with the target C at the middle of the editing window for maximum editing efficiency^16^ but not sgRNA5.2 which has the target C at the border of the editing window (Fig. 2A, 2B).

Gene editing outcomes in isolated clones revealed heterogenous genotypes with combinations of c.1392C>T and c.1393C>T in heterozygous or homozygous status. Since c.1392C>T was a synonymous SNV, 8 clones with heterozygous c.1393C>T from 20 analyzed clones were considered positive clones (Fig. 2C, 2D). However, in the clone CNM5-B, one extra heterozygous SNV c.1385C>A outside of the editing window was introduced (Fig. 2D), a rare 1 out of 20 event occurred during this editing. So only 7 positive clones were isolated with the correct genotype (Table 1). In general, although less specific BEs such as CBE4max_SpRY can provide options to avoid bystander editing, a more efficient and specific BE with adjusted sgRNA can significantly impact on-target editing outcomes.

### On-target efficiency highly depends on the target sequences

The optimized strategy was applied to all selected CNM mutations for CBE editing except one CNM mutation required an ABE. Although the 10 mutations are all for the CNM disease, editing efficiency varied from gene to gene and mutation to mutation (Fig. 2E, Table 1), similar to the base editing outcomes for random genes^16^. The *MTM1* mutations required more efforts to achieve on-target editing. We obtained only 6% editing efficiency (2 out of 33 clones) using CBE4max_SpG for the CNM-1 *MTM1* c.1456C>T, and no editing was achieved for the other two mutations (CNM-2 and CNM-3) with similar editing condition (Fig. 2E, Table 1 and supplementary Table S1). However, among the remaining 7 CNM mutations, only CNM-4 *DNM2* c.1105C>T could not be introduced to iPSCs with highly specific wild type BE4max and a highly specific sgRNA (supplementary Table S1). Even further increasing the concentrations of BE and sgRNA proved ineffective (data not shown).

Editing specificity was achieved in 5 successful generated iPSC CNM lines but not in 2 iPSC CNM lines (CNM-7 and CNM-9) (Fig. 2E, Table 1). For homozygous *RYR1* mutation, a low editing efficiency (∼14%) for CNM-9 c.8888T>C resulted in only iPSC clones with heterozygous SNVs but a high editing efficiency (∼45%) made successful editing for CNM-10 c.14545G>A (Fig. 2E, Table 1). On the other hand, for CNM-7 *BIN1* c.70C>T, a high 82% on-target editing was achieved with many clones carrying extra SNVs within the editing window (position 4 to 8) and outside of the editing window (position 1-2), indicating an expanded editing window (Fig. 2E, Table 1) caused by hyper BE activity. Moreover, clone CNM7-A with heterozygous *BIN1* c.70C>T also carried a 14bp insertion (c.69_82dup, p. Lys28ThrfsTer33) (supplementary Fig. S3A), an off-target due to single-strand break (SSB) by Cas9 nickase (nCas9) used in the current versions of BE^16^. Eventually for CNM-7, a heterozygous mutation was aimed to be introduced but only iPSC clones with homozygous SNVs showed no excess editing or insertion within the target sequence (supplementary Fig. S3). For other heterozygous CNM mutations such as CNM-6 *DNM2* c.1856C>T, a low editing efficiency (∼4%) was sufficient to generate 2 clones for this iPSC line.

Our results suggest a high editing efficiency is required to introduce homozygous mutations using BEs while a moderate editing efficiency can help reduce the screening size for clones with heterozygous mutations. Overall, the on-target efficiency and specificity highly depend on target sequences for each mutation and require optimization individually.

### Whole genome sequencing (WGS) verified on-targets and rare off-targets

WGS has been widely used for surveillance of gene editing activities.^17, 19^ Moreover, genetic alteration can also occur during iPSC culture^17^. To distinguish the genetic alterations associated with base editing from those gained from iPSC culture, we performed WGS for each generated iPSC CNM lines along with the parental iPSC line (supplementary Table S4) and developed a variant detection pipeline (Fig. 3A, see “Materials and Methods” for details). Clone-specific variants were identified by comparing the genome of each clone with the genome of the parental line.

**FIG 3.**
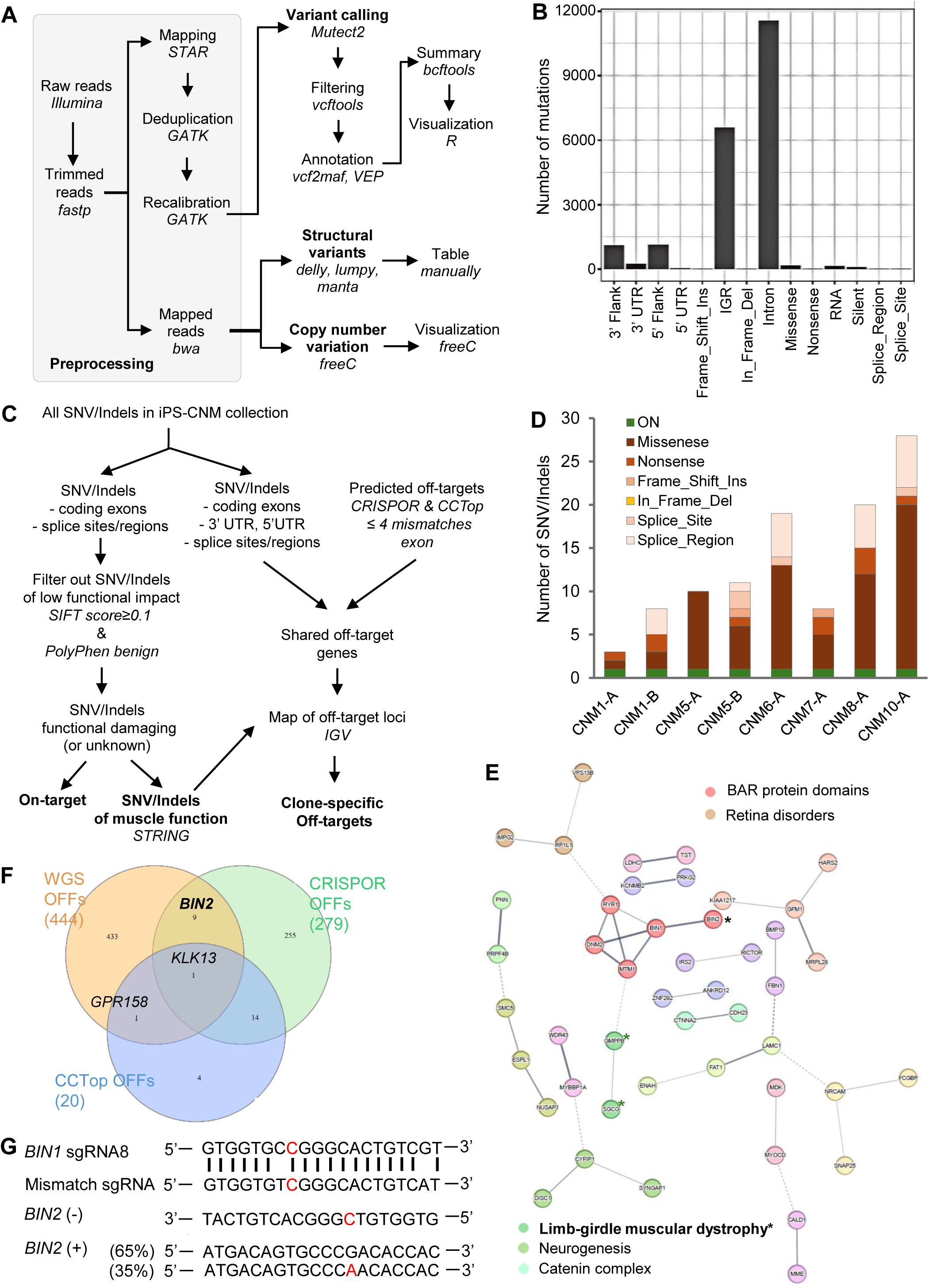
Quality control of the iPS-CNM collection using WGS. (A) A pipeline using WGS for genetic QC of generated iPSC lines. See detailed description from Methods section. (B) Types of SNV/Indels identified using Mutect2 from WGS data of 8 iPSC clones of 6 iPSC lines (supplementary Table S4) with reference to WGS data of unedited control line. (C) The on-target and off-target analysis pipeline starting from all SNV/Indels in the collection. (D) The refined coding and splice SNV/Indels for possible functional damaging ones including on-target (ON) using SIFT score and PolyPhen prediction (Fig.3C). The single on-target (ON) for each iPSC line was plotted in green, and different types of other SNV/Indels were plotted in orange series. (E) Perturbed proteins of refined SNV/Indels from Fig. 3D and their functional interaction network. The graph was generated using STRING and MCL clustering (find natural clusters based on the stochastic flow) was applied^38^. Highlighted clusters were showed in different colors. Proteins not in cluster (disconnected) were hidden from the graph. Green * marked two possible muscle proteins, and one black * marked the BIN2 as a member of BAR protein family. (F) Overlapped off-target genes (OFFs) from WGS analysis of possible coding and splice SNV/Indels (exons and splice site/region), with predicted OFFs of exon off-targets from CRISPOR and CCTop tool (maximum 4 mismatches). The Venn diagram was generated with InteractiVenn.^47^ (G) The *BIN2* off-target in clone CNM8-A due to only 2 mismatches of the sgRNA8 to *BIN2* target sequences. The mismatched sgRNA targeted the genomic minus (-) strand, leading to 35% conversion of A-to-G on the plus (+) strand.

Most SNV and small Indels were within the intergenic or intron regions (Fig. 3B). The possible damaging SNV/Indels including on-targets were filtered from SNV/Indels of coding exons and splice site/regions, by excluding those predicted to be low functional impact (Fig. 3C). We kept the SNV/Indels without functional predictions in the list for analysis (Supplementary Table S5). On-target SNVs were confirmed with WGS along with the correct zygosity, showing homozygous SNVs with 100% variant allele frequency (VAF) and heterozygous SNVs with roughly 50% VAF (Table 2, supplementary Fig. S4, S5). A variable number (3 to 28) of possible protein function damaging SNV/Indels for each clone were listed (Fig. 3D), but only two affected genes *GMPPB* and *SGCG* have a direct role in muscle function (Fig. 3E). However, the SNV of *GMPPB* occurred in the splice region without evidence of pathogenicity. The SNV of *SGCG* was heterozygous, but its related muscular dystrophy requires a homozygous mutation^40^. In addition, it is very unlikely that this mutation was introduced by base editing as the sgRNA6 and *SGCG* target region has more than 9 mismatches (supplementary Fig. S6A). Thus, no other function damaging SNV/Indels except the insertion *BIN1* c.69_82dup was further identified from the WGS variant list.

**Table 2.** Summary of the characterization on iPS-CNM collection.

| Cell Lines | Clones | On-target | WGS |  |  | Pluripotency |  |  | Usage |  |
| --- | --- | --- | --- | --- | --- | --- | --- | --- | --- | --- |
|  |  |  | Variant allele frequency | Off-targets | Other SNVs | Surface marker | Nuclear marker | Trilineage differentiation | Disease modeling | Gene therapy development |
| CNM1 | CNM1-A | *MTM1* c.1456C>T | 100% | - | - | - | - | - | Yes | Yes |
|  | CNM1-B | *MTM1* c.1456C>T | 100% | - | - | Yes | Yes | Yes | Yes | Yes |
| CNM5 | CNM5-A | *DNM2* c.1393C>T;<br>*DNM2* c.1392C>T | 42% | *KLK13* c.-5G>A, hom.;<br>*KLK13* c.-6G>A, hom.;<br>*KLK13* c.-12G>A, het. |  | - | - | - | Yes | No |
|  | CNM5-B | *DNM2* c.1393C>T | 56% | *KLK13* c.-5G>A, hom.;<br>*KLK13* c.-6G>A, het | *DNM2* c.1385C>A, het. | Yes | Yes | - | No | No |
| CNM6 | CNM6-A | *DNM2* c.1856C>T | 55% | - | - | Yes | Yes | Yes | Yes | Yes |
| CNM7 | CNM7-A | *BIN1* c.70C>T | 43% | *BIN1* c.69_82dup, het. |  | Yes | Yes | Yes | No | No |
|  | CNM7-B | *BIN1* c.70C>T | ~100% | - | - | - | - | - | No | Yes |
| CNM8 | CNM8-A | *BIN1* c.461G>A;<br>*BIN1* c.462G>A | 100% | *BIN2* c.458G>A, het. |  | Yes | Yes | Yes | Yes | No |
| CNM9 | CNM9-A | *RYR1* c.8888T>C | ~50% | - | - | - | - | - | No | Yes |
| CNM10 | CNM10-A | *RYR1* c.14545G>A | 97% |  |  | Yes | Yes | Yes | Yes | Yes |
| iPSC ctrl | Unedited parental | - | - | - | - | Yes | Yes | Yes | Yes | Yes |

The variant allele frequency was calculated based on allele counts from WGS data., except for CNM7-B and CNM9-A which was estimated from Sanger sequencing.

Off-targets are a critical concern for safety of using BEs^19, 20^. To avoid missing any important off-targets from our WGS analysis pipeline, we also predicted further exon off-targets by CRISPOR^22^ and CCTop tool^39^ (maximum 4 mismatches) (Fig. 3F, supplementary Table S6). Through comparing SNV/Indels of exons and splice sites/regions from WGS detection with predicted exon off-targets (Fig. 3C), one shared common off-target in the 5’UTR of *KLK13* was identified which occurred in both clones of the iPSC CNM-5 line (supplementary Fig. S6B). Another shared off-target gene between CCTop prediction and WGS was *GPR158* but the predicted off-target did not match the SNVs detected by WGS mutation analysis (supplementary Fig. S6C). Among 9 overlapping off-target genes identified by CRISPOR and WGS (Fig. 3F), only the gene *BIN2* showed the off-target with 2 mismatches to sgRNA8 in the corresponding cell line (Fig. 3G), while predicted off-targets from other genes could not be verified in the corresponding cell lines (supplementary Table S6). Thus, only off-targets in the 5‘UTR of *KLK13* gene occurred in iPSC CNM-5 line and a missense mutation in *BIN2* was in introduced to iPSC CNM8 lines (Table 2). Both genes *KLK13* and *BIN2* have no direction function in muscle.^41, 42^

Our results confirmed that off-targets occurred rarely in iPSC lines generated by base editing. Except rare events such as the small insertion in *BIN1* and one extra *DNM2* mutation, no further muscle disease causing SNV/Indels despite the on-target was detected in each iPSC CNM line using WGS.

### Structural variants (SVs) were barely induced by base editing in iPSCs

Our analysis revealed that no large copy number variations were detected using Control-FREEC, a tool for detection of copy-number changes and allelic imbalances.^32^ Not surprisingly, a number of SVs such as inversion, deletion, translocation and duplication were detected using different SV caller DELLY^33^, LUMPY^34^, and (1.6.0)^35^. Only 1 to 2 SVs for each clone detected by at least two SV callers were considered moderately confident (supplementary Table S7). No common SVs were detected within clones of one iPSC CNM line or within different iPSC CNM lines. Moreover, the SVs did not occur near the base editing target region (within 200bp) (supplementary Fig. S4-S5), which strongly suggests that the changes were not caused by base editing.

Therefore, we can conclude that the base editing process had minimal impact on the global genome stability without inducing significant SVs in generated iPSC CNM lines.

### Pluripotency validation for iPSC CNM lines

Assessing the pluripotency of generated iPSC CNM lines are critical for downstream analysis. A comprehensive characterization was conducted using well established protocols.^26^ We observed some positive edited clones lost iPSC morphology (data not shown) therefore were initially excluded from expansion and quality controls. Only iPSC clones with validated on-target editing and good iPSC morphology were further proceeded to pluripotency assessment. All tested clones had good quality (Table 2), showing positive signals of nuclear markers OCT4, SOX2 and NANOG (Fig. 4A and supplementary Fig. S7), almost 100% cells with surface pluripotency markers (SSEA-4, TRA-1-60, TRA-1-81) and low amount cells (<3%) with differentiation marker SSEA-1 (Fig. 4B, 4C). In addition, iPSCs were able to turn into cells from all three embryonic germ layers (Fig. 4D).

**FIG 4.**
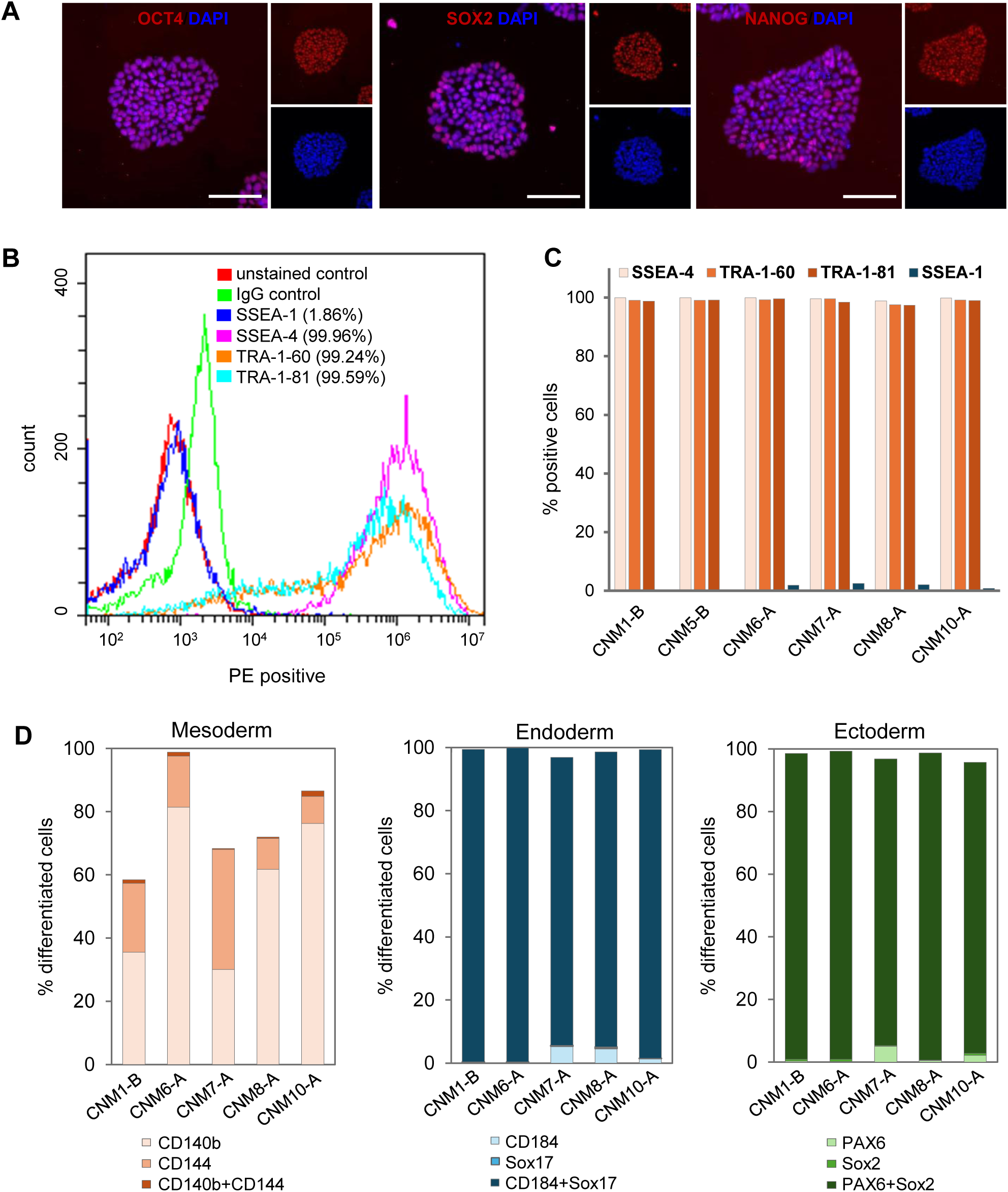
Characterization of iPSC pluripotency for CNM lines. (A) Immunostaining of CNM6-A showing merged overlays (purple) of pluripotency markers OCT4, SOX2 and NANOG (red) and DAPI counterstaining (blue). Scale bar = 100 µm. (B) Flow cytometry analysis of CNM6-A for pluripotency markers SSEA-4, TRA-1-60, TRA-1-81 and differentiation marker SSEA-1. Unstained and IgG isotype controls were included. (C) Summary of the pluripotency analysis using surface markers for selected CNM lines. The percentage of positive cells for each surface marker was detected using flow cytometry as showed in Fig. 4B and summarized here. (D) Trilineage differentiation efficiency was shown as percentage of differentiated cells per germ layer. Marker expression was measured by flow cytometry based on single- and double-positive populations.

The iPSC clones for iPSC CNM lines passed through genetic and pluripotency QCs meet the criteria for downstream applications as proposed in Table 2.

## Discussion

Establishing a reliable genotype-phenotype correlation for neuromuscular disorders such as CNM is difficult due to a unique genomic or epigenetic background of individual patient, which can severely mask the primary effects of the disease-causing mutations. To overcome this limitation, we generated a collection of iPSC lines carrying patient-specific CNM mutations with one parental iPSC line using base editing. This will allow direct investigation of single mutation impact on muscle functions at protein, cell or tissue level in a relatively similar background, in order to gain genotype-phenotype correlation and to seek precise treatment.

We applied different combinations of BEs and sgRNAs to generate X-linked, heterozygous or homozygous mutations of 4 CNM genes. Overall editing results confirmed that the general editing efficiency of BEs from highest to lowest is BE4max, CBE4max_SpG, CBE4max_SpRY^16^. During this process, we observed variable editing efficiencies among CNM genes or mutations of the same CNM gene and also failure of editing, which highly depend on a number of factors such as local sequence context, target position within the editing window, chromatin accessibility and sgRNA binding efficiency.^43^

For iPSC line generation, a small number of edited, high quality iPSC clones were sufficient for downstream studies.^44^ Therefore, we learned that iPSC line generation using BEs required different strategies for introducing heterozygous and homozygous mutations. For heterozygous mutation introduction, a low to moderate editing was sufficient to obtain several edited clones, but high editing efficiency often created a large number of clones with homozygous SNVs. Moreover, there is higher risk of introducing unintended changes into neighboring bases due to an expanded editing window because the hyperactive BEs retained high catalytic activity across a larger portion of the target region.^45^ In contrast, introduction of homozygous mutations requires higher BE activity as we encountered difficulties of using a less efficient ABE8e_SpRY to obtain iPSC clones with a homozygous mutation. A suitable editing efficiency by titrating the expression level or the target exposure times of BEs from plasmid or mRNA^25^ can further improve our iPSC line generation system.

The BE4max and its variants use a Cas9 nickase (nCas9) which cuts the opposite strand when deamination occurs on the other strand^16^. No DSBs reduce the risk of large deletions, insertions, or random chromosome mixing. Our WGS analysis pipeline detected no copy number variations and only a few SVs which were not within 200bp of the target, suggesting these SVs are not caused by base editing but likely occur during iPSC expansion^26^. However, although relatively rare, we did detect a small insertion in generating iPSC CNM-7 line due to the nCas9 activity because the targeted base modification and the opposite-strand nick are processed as a functional DSB followed by DSB repair.^46^

Off-targeting poses a major safety risk associated with base editing, which occurs when the sgRNA binds to unintended genomic loci that share sequence homology with the target protospacer.^19, 20^ Although there are BE-specific methods such as CHANGE-seq-BE for detecting off-targets^20^, we showed WGS can cover more genetic QC measurements from detection of BE introduced on-targets and off-targets, to identification of possible pathogenic SNV/Indels and large SVs emerged during iPSC expansion. Unlike the whole edited cell pool containing a large number of off-targets with low VAF, individual iPSC clone derived from that pool only inherits a small fraction of those off-targets. The clone-specific off-targets typically appear in heterozygous or homozygous status so they are easy to be detected via WGS analysis due to their high VAF. We also showed only few overlapping between predicted off-targets by tools such as CRISPOR^22^ or CCTop^39^ for bulk editing events and WGS detected clone-specific variants. The WGS was sufficiently sensitive to detect the small insertion in *BIN1* gene in the case of CNM-7 or the predicted top off-target in *BIN2* gene in the case of CNM-8.

The advantage of using WGS for genetic QC is that it covers partially standard QC for iPSCs such as SNP array analysis or karyotyping. But G-banding is still necessary for balanced translocations and polyploidy. Moreover, pluripotency QC is critical for using the iPSC lines for downstream applications. Although we observed some differentiated clones after gene editing and further prolonged *in vitro* expansion (data not shown), clones with good iPSC morphology passed the standard pluripotency QC. Eventually, for each iPSC CNM lines, not all edited clones with good iPSC morphology went through full characterization. So for downstream application such as muscle modeling, we took at least one such fully characterized clones with additional clones as standard practice in iPSC disease modeling.^44^

There are a few of limitations to this study. We did not conduct multiple transfections for each base editing as we generally aimed for a couple edited good clones for each cell line. Therefore, our results on editing efficiency can only provide conclusion about general principles such as BE4max or CBE4max_SpG showed higher editing efficiency than CBE4max_SpRY. Additionally, only a small number of selected clones were selected for WGS due to cost reason. Therefore, the WGS pipeline can be further refined for accuracy, if WGS data from more clones become available. Two iPSC CNM lines (CNM-7 and CNM-9) in collection do not share the same zygosity as patients, but they can be used to screen BEs and sgRNAs for mutation correction prior to validating using precious patient-derived cells. An expanded iPS-CNM is expected through the inclusion of additional clones from more mutations, thereby providing a comprehensive understanding of CNM.

Despite this is a preliminary version of iPS-CNM, we generated 7 iPSC CNM lines using BEs of which 6 underwent full characterization. Most importantly, we built a WGS analysis pipeline to validate the on-targets, detect off-targets and sort out SVs. The iPSC CNM lines with fully described genetic background can serve as valuable material to conduct disease modeling or to screen BEs and sgRNAs for patient mutation correction. The WGS QC strategy can also be easily adapted to off-target surveillance for gene therapy development.

## Acknowledgments

We are grateful to Barbara Beuerle for technical support.

## Authors’ Contributions

H.W. designed experiments, analyzed data and wrote the manuscript. I.S. (Ida Steinkirchinger), V.H., H.K., and S.K., constructed, prepped, sequence verified sgRNA constructs. L.G. performed cell sorting. I.S. (Insa Staecker), A.E., and V.H, conducted transfection, colony picking, cryopreservation, and iPSC quality assurance. M.K., I.S. (Ida Steinkirchinger), and J.H., provided assistance with iPSC quality assurance. I.S. (Insa Staecker), A.E., I.S. (Ida Steinkirchinger), H.K., and H.W., conducted genotyping of edited clones using Sanger sequencing. L.S. provided support on the standard iPSC banking and quality controlling. C.P. performed NGS data processing and analysis. All authors reviewed the manuscript.

## Supplementary Material

Supplementary Figures

Supplementary Tables

