## Supplementary Figures for "iPS-CNM: an iPSC collection generated by base editing for centronuclear myopathy"

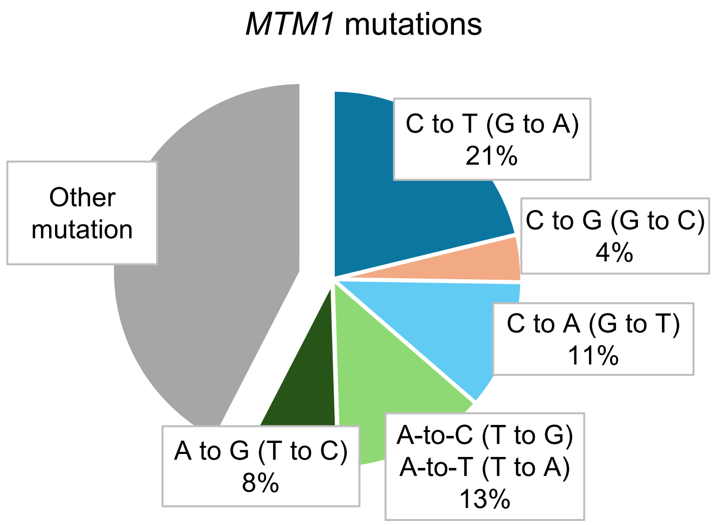


Fig. S1. Base changes of reported mutations in *MTM1* gene. The mutations included the pathogenic and likely pathogenic variants in ClinVar database (total 253 mutations until 2025).

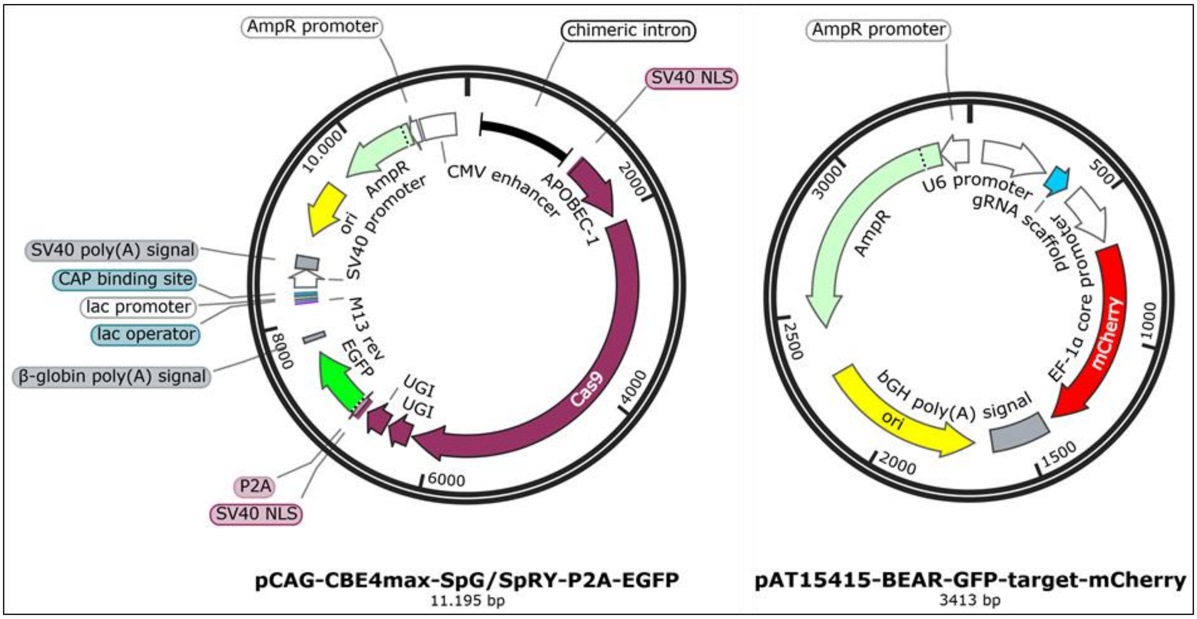


Fig. S2. Plasmid maps of base editors and sgRNA.

The pCAG-CBE4max-SpG-P2A-EGFP (Cat. No. Addgene: 139998) and pCAG-CBE4max-SpRY-P2A-EGFP (Cat. No. Addgene: 139999), sharing the identical vector backbone (left) with EGFP reporter, but included different Cas9 variants (either SpG or SpRY). Plasmid map pAT15415-BEAR-GFP-target-mCherry (right), carrying a mCherry reporter and a sgRNA scaffold, which was replaced with sgRNA sequences targeting CNM genes. Figure created with SnapGene and plasmid sequences were adapted from Addgene.

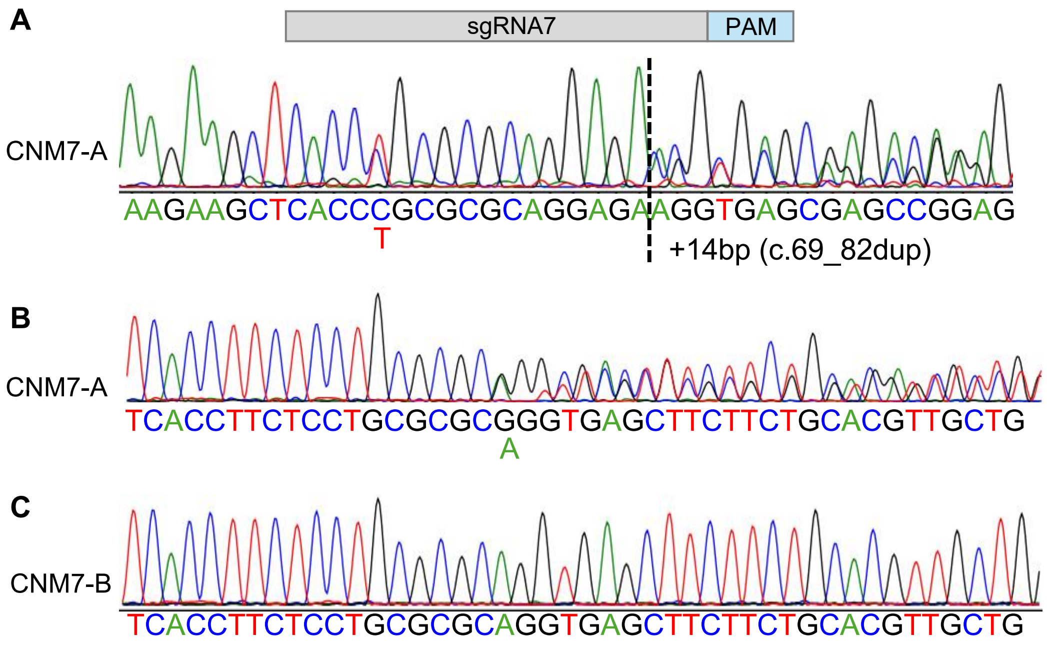


Fig. S3. Sanger sequencing for representative CNM7 clones. (A) The clone CNM7-A carried one heterozygous c.70C>T and a 14bp insertion (c.69_82dup) due the cleavage of the nCas9 of BE with forward sequencing of the target region. (B) The clone CNM7-A with reverse sequencing of the target region (C) The representative clone carried homozygous c.70C>T without additional SNV in the target gene CNM7-B.

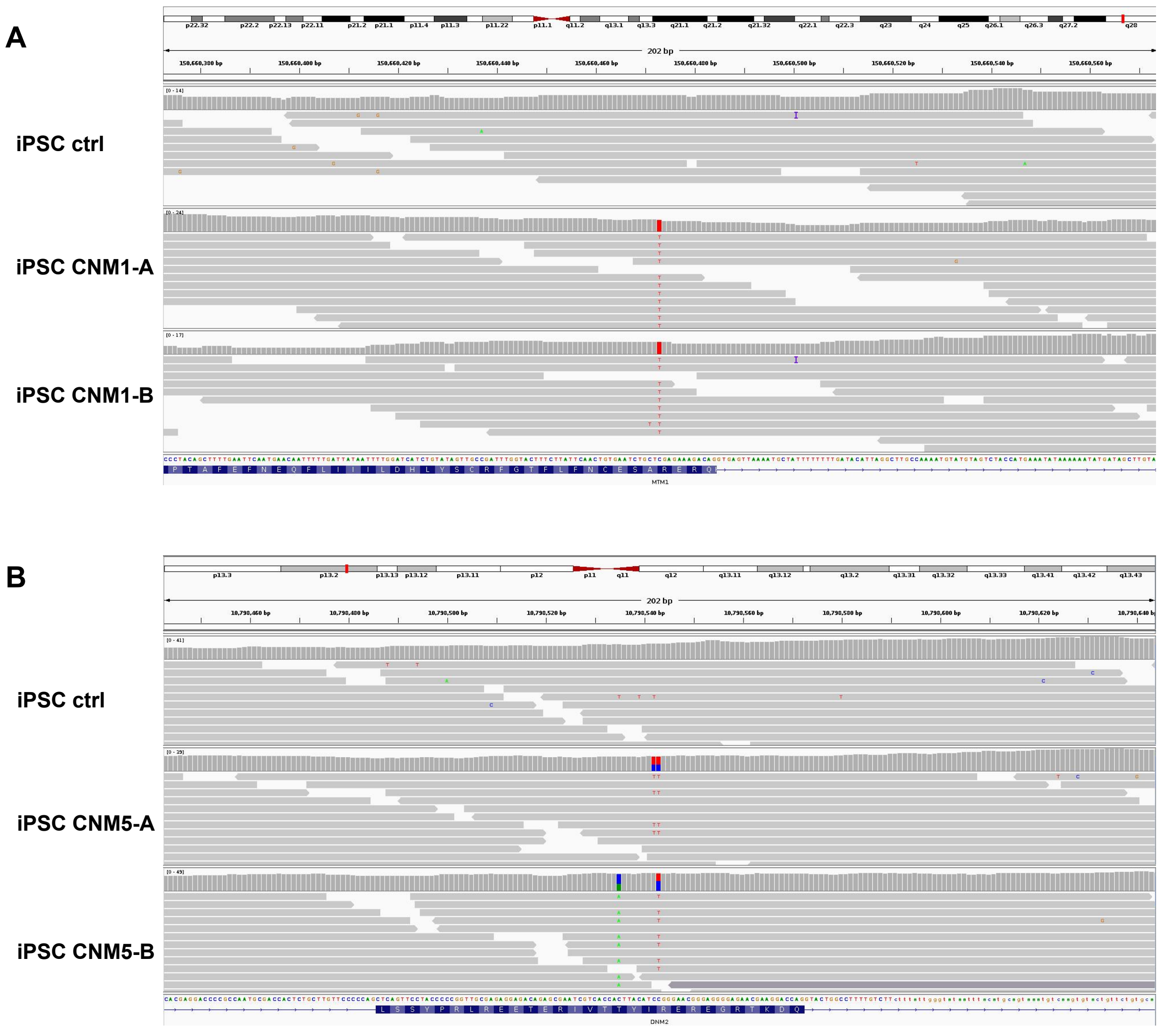


Fig. S4. On-targets of CNM SNVs detected by WGS for CNM1 and CNM5. The on-target visualization in IGV browser for the single target and a surrounding genomic region of 200bp for each target with sequencing coverage and genomic loci. The sequence was aligned to the parental unedited line (iPSC ctrl). (A) Two clones for CNM1 iPSC lines with on-target editing in *MTM1* c.1456C>T on the X chromosome. (B) Two clones for CNM5 iPSC lines with on-target editing in *DNM2* c.1393C>T (heterozygous). Clone CNM5-A carried a synonymous bystander editing c.1392C>T (heterozygous). Clone CNM5-B carried a heterozygous SNV c.1385C>A.

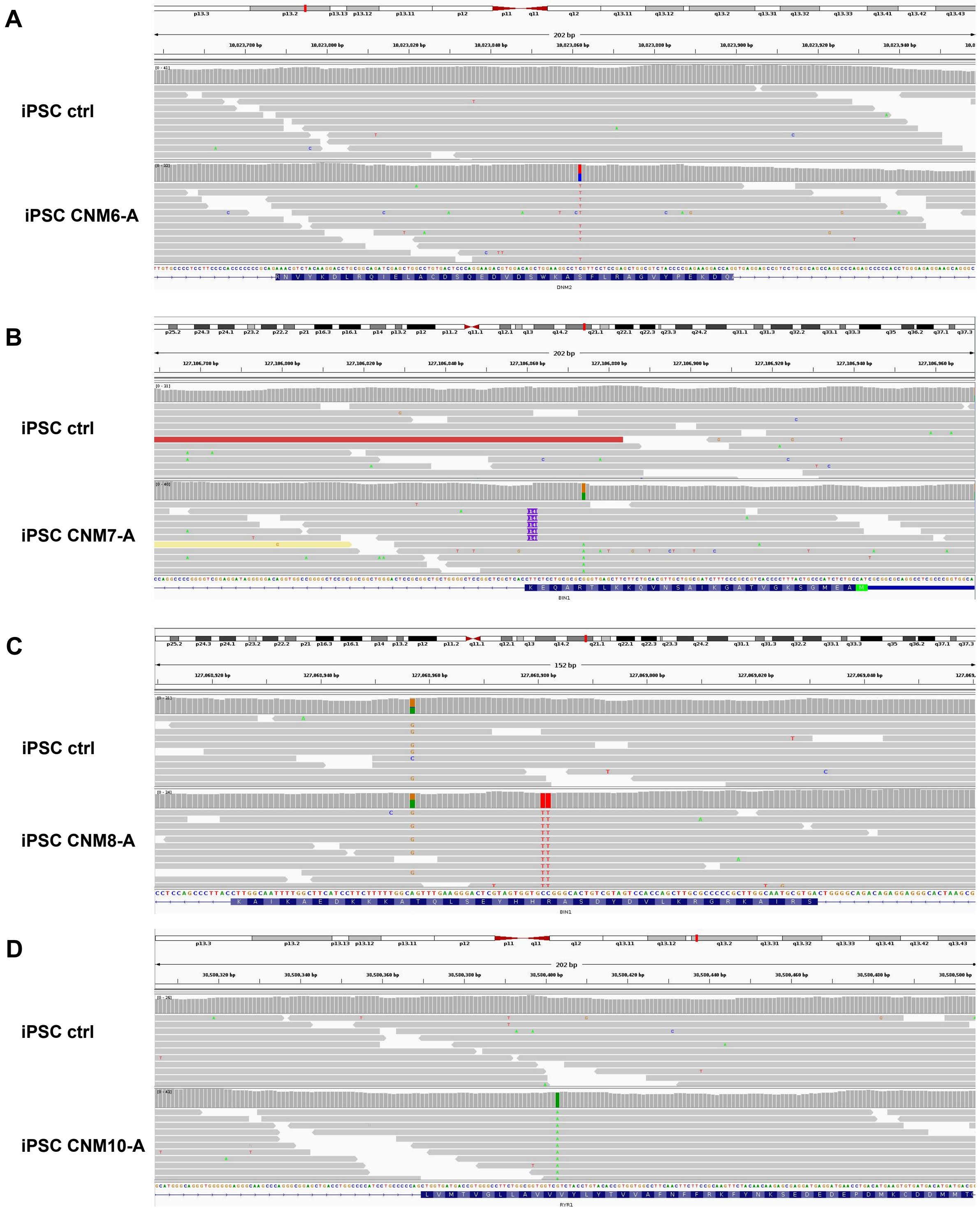


Fig. S5. On-targets of CNM SNVs detected by WGS for CNM6, CNM7, CNM8 and CNM10. The on-target visualization in IGV browser for the single target and a surrounding genomic region of 200bp for each target with sequencing coverage and genomic loci. (A) WGS confirmed the heterzygous on-target SNV with ~50% allele frequency in CNM6-A iPSC line. (B) WGS confirmed the 14bp insertion caused by BE in CNM7-A iPSC line. (C) WGS confirmed the homozygous on-target SNV with a bystander editing in CNM8-A iPSC line. (D) WGS confirmed the homozygous on-target SNV in CNM10.

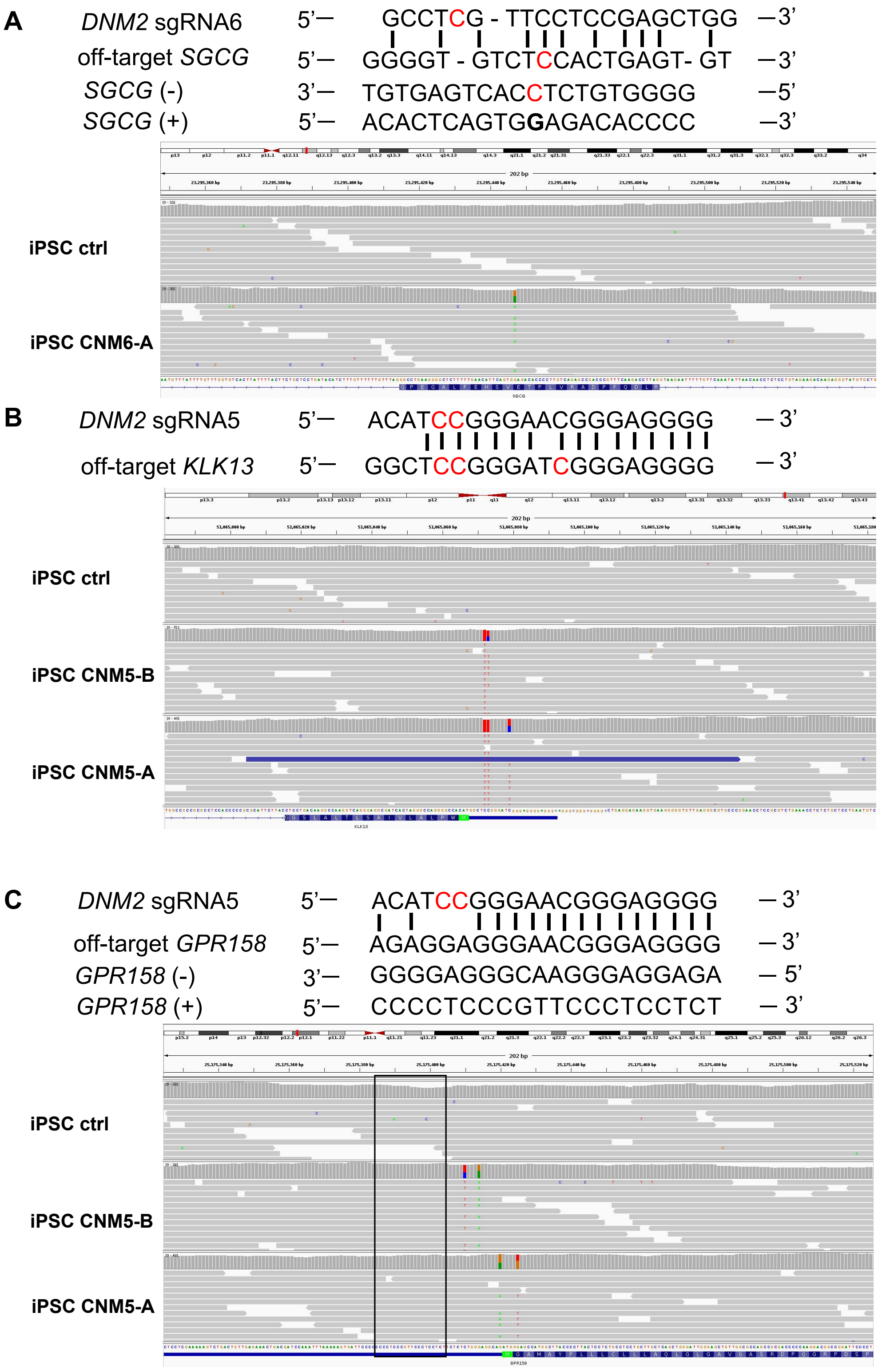


Fig. S6. SNVs as off-targets or not off-targets. (A) The *SGCG* SNV in CNM-6. SgRNA6 and the *SGCG* region with SNV had 9 mismatches. (B) The *KLK13* in CNM-5. SgRNA5 and the target region had 4 mismatches as predicted by CRISPOR. The off-targets were detected in both CNM-5 clones by WGS. (C) SgRNA5 and the predicted *GPR158* off-target region (within the black frame) had 4 mismatches as predicted by CCTop but no SNV was identified in the predicted region by WGS. The regions with detected SNVs by WGS had poor alignement with sgRNA5 therefore these SNVs were not off-targets.

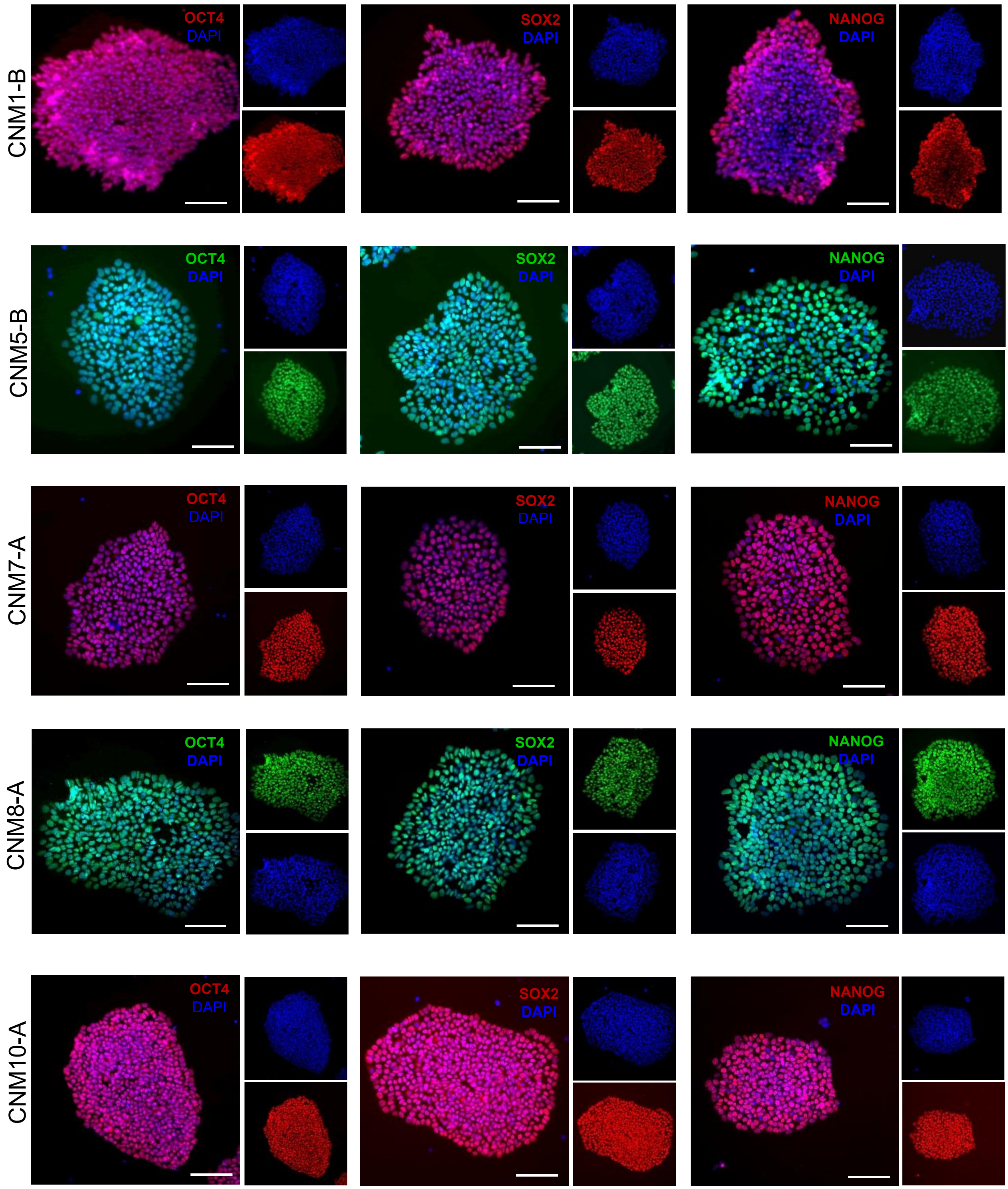


Fig. S7. Staining of CNM iPSC lines with pluripotency markers OCT4, SOX2 and NANOG (red or green as indicated with different secondary antibodies) and DAPI counterstaining (blue). This is the extended data of Fig. 4A. Scale bar = 100 µm.
